# The distribution of integer partitions in human genome-wide genealogies reflects gene flow from a super-archaic lineage

**DOI:** 10.64898/2026.07.14.738421

**Authors:** Alexander Mackintosh

## Abstract

Several recent studies have found evidence for ancient gene flow between the ancestors of modern humans and an unsampled super-archaic lineage. Here we present a new, simple approach for characterising this process given genome-wide genealogies sampled from a single population. We summarise genealogies as distributions of integer partitions and show that this captures the temporal signal of tree imbalance left by ancient gene flow. We analyse genealogies from modern humans and find that the integer partition distributions are inconsistent with a history of panmixia but can be explained by gene flow from a super-archaic lineage. Our analysis favours a model of continuous gene flow over pulse-admixture and also recovers a bottleneck in the ancestors of modern humans. This work highlights a clear signal of ancient structure in genealogies of modern humans and provides an inference approach that complements existing methods.

## Introduction

Several recent studies have found evidence for ancient gene flow between the ancestors of modern humans and a “super-archaic” hominin lineage (Fan *et al*. 2023; Ragsdale *et al*. 2023; Cousins *et al*. 2025; Zhang *et al*. 2026; Loya *et al*. 2026; Rogers *et al*. 2026). The super-archaic lineage is estimated to have diverged from modern humans at least 1 mya and there is speculation that it corresponds to *Homo erectus*. Reconstructing the process of gene flow between these groups is a technical challenge given the lack of genetic data for any super-archaic individual. Recent studies have approached this problem by leveraging patterns of linkage disequilibrium in genome sequences of modern humans (Ragsdale *et al*. 2023; Cousins *et al*. 2025; Zhang *et al*. 2026) or exploiting the effect of super-archaic gene flow on genealogical relationships between modern humans, Neanderthals and Denisovans (Loya *et al*. 2026; Rogers *et al*. 2026). These alternative approaches have produced broadly consistent results, but there is still uncertainty about the timing, magnitude and duration of gene flow.

Here we investigate an alternative source of information for detecting and characterising ancient gene flow - the distribution of integer partitions of a genealogy. The remainder of this article is structured as follows. First, we briefly explain the distribution of integer partitions and its connection to other properties of genealogies. Next, we show that a history of ancient admixture and a history of panmixia with equivalent rates of pairwise coalescence through time have distinct integer partition distributions. Finally, we calculate these distributions from genome-wide genealogies of modern-day humans and show that they are best explained by continuous gene flow from a super-archaic lineage.

## Results

### The distribution of integer partitions of a genealogy

As an example, consider the coalescent process for *n* = 4 lineages that have been sampled from a single population (Figure 1A). At the time of sampling there are four exchangeable lineages, each with one descendant, so the integer partition of the sample at this time is (1, 1, 1, 1). Moving backwards in time, the first coalescence event results in a new partition – (2, 1, 1) – as there are three lineages, one of which has two descendants. The second coalescence event results in either the partition (3, 1) or (2, 2), reflecting the two possible unlabelled topologies for a sample of *n* = 4. The final coalescence event in the history results in a partition of (4). Any marginal genealogy has exactly one unique succession of integer partition states (Figure 1A), yet the distribution of integer partitions across many genealogies reflects both topology and rate of coalescence in the underlying ancestral recombination graph (Figure 1B and C).

**Figure 1:**
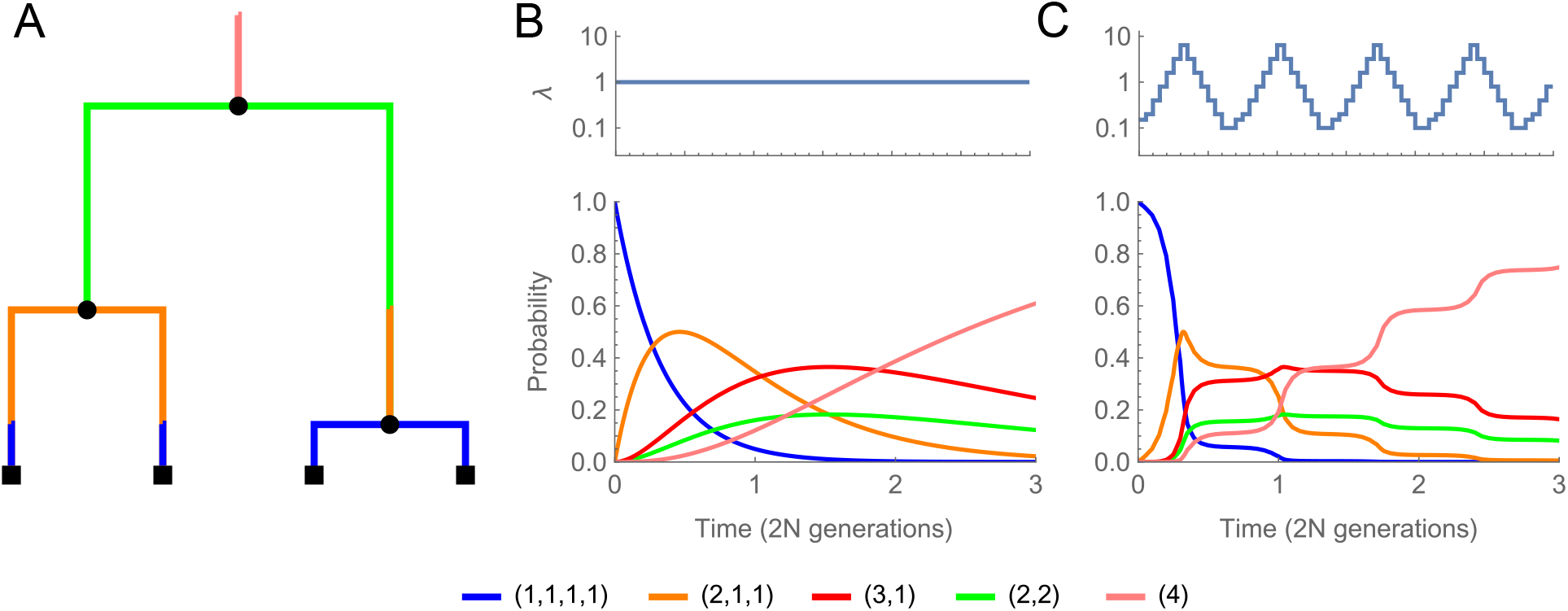
The distribution of integer partitions for a sample of four lineages from a panmictic population. **(A):** An example genealogy where branches are coloured by integer partition state. **(B):** The probability distribution for all five possible partitions under constant population size with time measured in units of 2*N* generations. **(C):** The probability distribution for a population of variable effective size, where the rate of coalescence relative to 1*/*2*N* (*λ*) oscillates through time.

Integer partitions appear in calculations for obtaining properties of genealogies under the coalescent (Pitman 1999; Rogers 2022; Lohse *et al*. 2016; Sendrowski and Hobolth 2026), but have received little attention as a summary for statistical inference. They have a direct connection to the site frequency spectrum (SFS); while the SFS represents the expected total length of branches with *i* descendants in a genealogy, distributions of integer partitions track the co-occurrence of these branch-types through time. Although here we only consider the marginal distribution of each integer partition (Figure 1), their joint distribution is in fact a lossless summary of the information contained in genealogical trees with exchangeable lineages.

The probability of observing a specific partition at time *t* can be calculated in closed form or numerically using matrix exponentiation (see Methods). The numerical approach naturally lends itself to calculations across epochs with piece-wise constant rates (Figure 1C). Discrete pulses of admixture and continuous gene flow can be accommodated in this framework by keeping track of how the partitions are distributed across populations, which we make use of in the next two sections.

### Distinguishing between panmixia and ancient structure

We next demonstrate that the distribution of integer partitions for a single population contains information about past gene flow. We focus on a simple model of pulse-admixture involving a focal population (A) and a unsampled population (B). The parameters of the model are inspired by the results of Cousins *et al*. (2025) (see the caption of Figure 2 for exact values). We generate a panmictic analogue of this model by calculating the expected rate of coalescence in narrow epochs for a pair of lineages sampled from population A. Figure 2 shows the distribution of integer partitions for the admixture and panmictic models between the time of admixture (*T*_0_) and the divergence time of the admixing populations (*T*_1_), conditioned on *n* exchangeable lineages being present at *T*_0_. For *n* = 2 the distributions are equivalent to the expected pairwise coalescence CDF and, as a result, the admixture and panmictic models have identical distributions. For *n ≥* 3, the distributions show differences that are expected consequences of admixture. For example, when *n* = 4 the admixture distribution has a scarcity of (2, 2) partitions relative to panmixia. This is because the (2, 2) partition cannot be generated between *T*_0_ and *T*_1_ if at time *T*_0_ three lineages move into one population and one lineage moves into the other. Similar logic applies to the results for *n* = 5, where some partitions have identical distributions under panmixia (e.g. (4, 1) and (3, 2)) but admixture generates asymmetry in their frequency. While unbalanced trees are the clearest signal of admixture for *n* = 5, the effect of admixture can still be observed for *n* = 3 despite there only being one possible topology (Figure 2). This is because the distribution of integer partitions contains information about the variance in coalescence rate across loci due to admixture. Together these results show that ancient gene flow can be detected given sufficient knowledge of the distribution of integer partitions.

**Figure 2:**
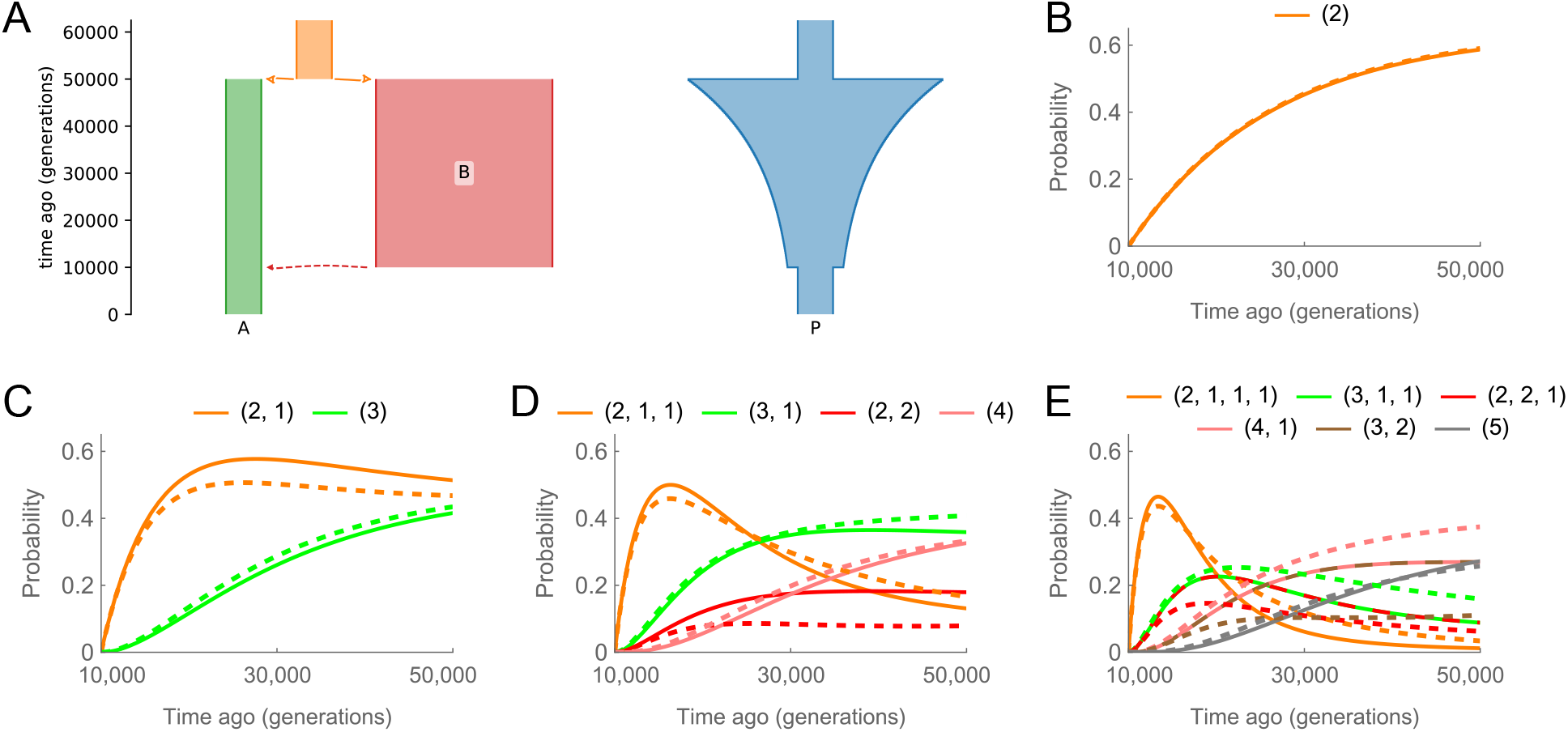
Integer partition distributions under panmixia and admixture. **(A):** A simplified demographic history of pulse-admixture based on the results of Cousins *et al*. (2025). Population A (*N*_*e*_ = 8 *×* 10^3^) receives a pulse of admixture from population B (*N*_*e*_ = 4 *×* 10^4^) at *T*_0_ = 1 *×* 10^4^. This results in 21% of lineages sampled from population A tracing their ancestry back to population B. These populations diverged at *T*_1_ = 5 *×* 10^4^. A panmictic demographic model which has the same pairwise coalescence rate is shown to the right. **(B-E):** Distributions of integer partitions conditional on *n* = 2 **(B)**, *n* = 3 **(C)**, *n* = 4 **(D)** or *n* = 5 **(E)** lineages being present at *T*_0_. In each plot the solid line shows the expectation for the panmictic model, while the dashed line shows the expectation under pulse-admixture. Distributions for the first partition (e.g. (1, 1, 1)) are not shown to maximise clarity.

### Evidence for ancient gene flow in human genome-wide genealogies

Integer partitions cannot be directly observed from genotype data. They are nonetheless reflected in patterns of linked polymorphisms and are accessible in reconstructed genome-wide genealogies.

Deng et al. have recently developed efficient methods for MCMC sampling genome-wide genealogies (SINGER; Deng *et al*. 2025b) and re-estimating their node ages without a coalescent prior (POLE-GON; Deng *et al*. 2025a). Here we re-analyse genealogies that were estimated for modern humans using these methods (YRI population of the 1000 genomes project; Byrska-Bishop *et al*. 2022; Deng *et al*. 2025a). We record the integer partition distribution for a sample of *n* = 5 between 8,000 generations ago (*∼* 220 kya assuming a generation time of 28 years) and 70,000 generations ago (*∼* 2 mya). We fit models to all seven integer partitions for a sample *n* = 5, but we pay particular attention to the four partitions that reflect topology and are therefore the most informative about past gene flow (Figure 3). Because inferred genealogies are imperfect reconstructions (Brandt *et al*. 2022; Martin 2026), we also apply a bias correction step to the model fitting procedure where bias is estimated by comparing integer partition distributions in true genealogies and those inferred with SINGER and POLEGON (see Methods).

**Figure 3:**
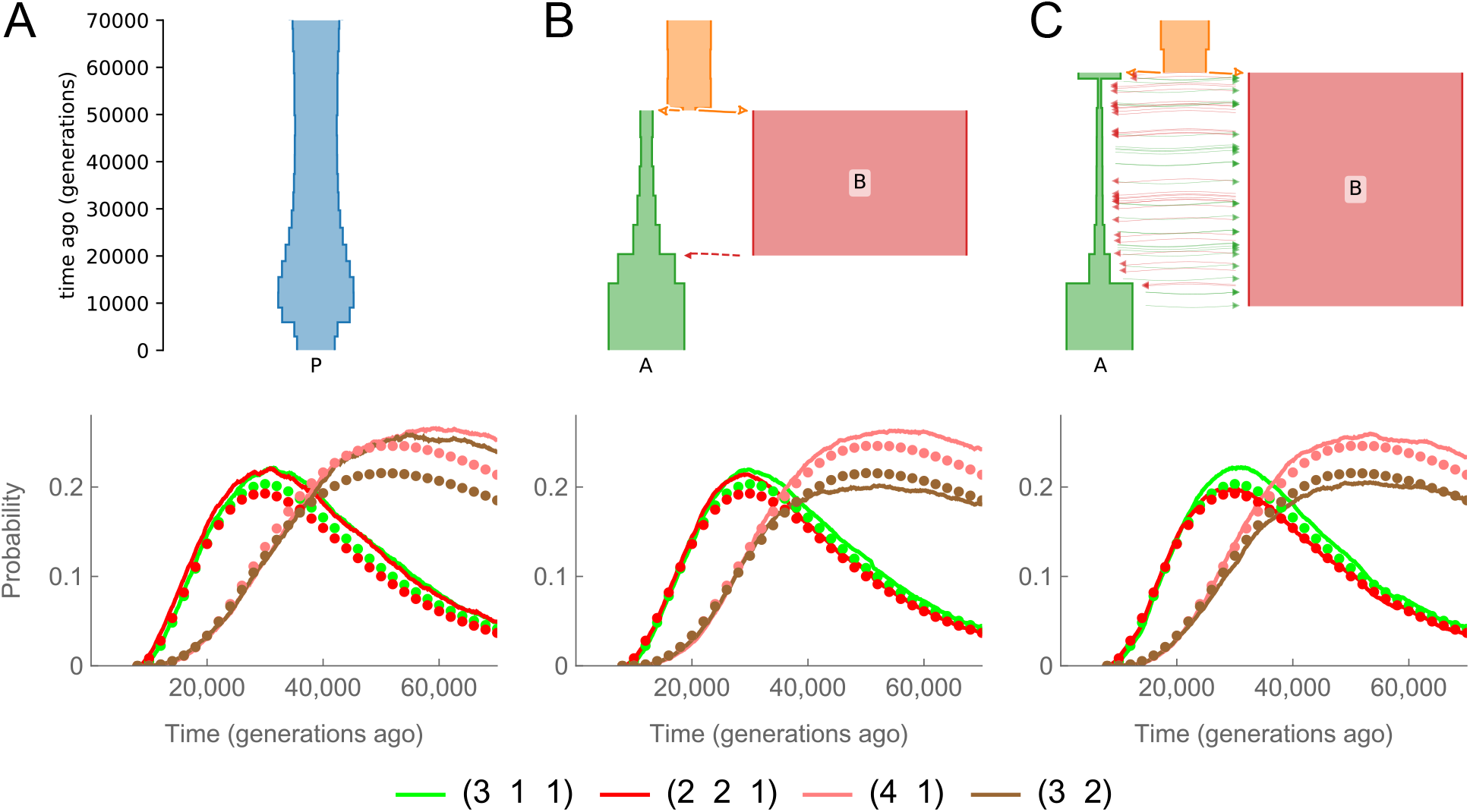
Distributions of topologically informative integer partitions for alternative models of population history. **A:** A panmictic history estimated from pairwise coalescence times in human genealogies. **B:** A model of pulse-admixture estimated from integer partition distributions in human genealogies. **C:** A model of continuous gene flow estimated from the same data. The lines in the lower plots show the bias-corrected analytic prediction of each model whereas the points show the distributions from human genealogies.

The distributions from human genealogies show a clear excess of (4, 1) partitions over (3, 2) in deep time and a more subtle excess of (3, 1, 1) partitions over (2, 2, 1) (Figure 3). These partitions are expected to have equivalent distributions under panmixia, suggesting that the human data is inconsistent with such a history. To test this, we estimated a piece-wise constant *N*_*e*_ trajectory from the distribution of pairwise coalescence times in the human genealogies (Figure 3A) and calculated the expected integer partition distributions under this history while accounting for the bias introduced by genealogy reconstruction. This model is indeed a poor fit to the data as it cannot explain the imbalance between integer partitions with an equal number of lineages (Figure 3A).

We next fit a model of pulse-admixture to the empirical integer partition distributions, while again accounting for reconstruction bias. Here population A contributes the majority share of ancestry to modern humans and receives a pulse of admixture from population B at time *T*_0_ (Figure 3B). This model predicts tree imbalance in deep time and provides a better fit to the data (Figure 3B; increase in composite likelihood of Δ*lnCL* = 8.06 over panmixia). The admixture timing was estimated to be 2.0 *×* 10^4^ generations ago (*∼* 560 kya) with a minor admixture proportion of *α* = 0.16. The divergence time between populations A and B was estimated at 5.1*×* 10^4^ generations ago (*∼* 1.4 mya). Our estimates of *N*_*e*_ in population A are across ten epochs (Figure 3B), whereas we set the *N*_*e*_ of population B to be constant through time. Our estimate of *N*_*e*_ in population B (1 *×* 10^5^) is the maximum value allowed in our model fitting. We also fit a model with continuous bidirectional gene flow between populations A and B (Figure 3C). We estimate that migration at rate *m*_*e*_ = 7.9 *×* 10^*−*5^ begins 9.3 *×* 10^3^ generations ago (*∼* 260 kya) and continues into the past until the populations merge 5.9 *×* 10^4^ generations ago (*∼* 1.6 mya). This model has the same number of parameters as the pulse-admixture model but provides a slightly better fit to the data (Figure 3C; Δ*lnCL* = 0.70).

Our analysis suggests that unbalanced integer partitions in deep time are explained by gene flow from a super-archaic lineage. We performed additional analyses to investigate whether there are other plausible explanations for this result. First, we simulated a panmictic population with hard selective sweeps. We set the waiting time between sweeps to be 6.4 *×* 10^4^ generations per-Mb, resulting in a genome-wide rate of one sweep every 20 generations. These simulations do lead to unbalanced integer partitions (Figure S1), but the effect is much weaker than what we observe in the human genealogies. We also simulated a subdivided population consisting of many demes. Specifically, we simulated a stepping-stone model with 30 demes of effective size *N*_*e*_ = 1000 and symmetrical migration between adjacent demes at rate *m*_*e*_ = 1 *×* 10^*−*3^. This simulation generates strong asymmetry between integer partitions (Figure S1). However, the asymmetry favours balanced trees, which is the opposite of the pattern we observe in human genealogies. These analyses suggest that other selective and demographic processes will affect our results to some degree, but that gene flow from a super-archaic lineage remains the best explanation for the observed data.

## Discussion

Here we have presented a novel approach for detecting and characterising ancient gene flow. Our summary of genealogies – distributions of integer partitions – capture two effects of gene flow on genealogical trees; variation in coalescence rate across loci and tree imbalance (Kirkpatrick and Slatkin 1993; Durand *et al*. 2011; Li and Wiehe 2013; Dilber and Terhorst 2022). The most similar existing approach to ours is that of Pope *et al*. (2023), who use rates of coalescence in labelled samples of *n* = 3 to infer multi-population demographic histories from reconstructed genealogies. While an unlabelled sample of *n* = 3 does contain some information about past gene flow, our results suggest that the topology information present at *n >* 3 provides a clearer signal (Figure 2 and 3). Two other recent studies have used genome-wide genealogies to characterise ancient gene flow in humans. Zhang *et al*. (2026) exploit covariance in lineage persistence and genomic span to detect admixture tracts, whereas Loya *et al*. (2026) fit a mixture model of ancestry using pairwise coalescence rates between modern and archaic humans. The main signal of gene flow that we have focused on – tree imbalance – is not used by either of these methods. In fact, our approach has more in common with SFS-based analyses that implicitly use information about tree topology (Durvasula and Sankararaman 2020; Fan *et al*. 2023). The main difference is that the SFS only contains information about expected branch lengths, whereas our summary captures information about the joint distribution of different branch types.

Our main result is that integer partitions in human genome-wide genealogies cannot be explained by a history of panmixia. This result does not require any model fitting as the imbalance in topology is clearly visible in deep time (Figure 3) and a panmictic population history cannot recover this. Model fitting was nonetheless required to show that realistic scenarios of super-archaic gene flow can recover this feature. Although the specific models we fit are more illustrative than precise reconstructions of human history, the analysis does provide some useful insights. For instance, the continuous gene flow model is favoured over a scenario of pulse-admixture, consistent with the results of Ragsdale *et al*. (2023). Additionally, we infer a bottleneck in population A with a gradual recovery, which was also found by Cousins *et al*. (2025). Our analysis confirms that this bottleneck is not an artifact of assuming a model of pulse-admixture, as it is still inferred under a scenario of continuous gene flow.

Our approach for inferring ancient gene flow does have several limitations. First, it is sensitive to evolutionary forces other than gene flow that lead to variance in the rate of coalescence across the genome. In our analysis of human genealogies we sampled loci which experience similar effects of background selection (see Methods), yet our results could still be influenced by old selective sweeps and balancing selection. Second, it requires that inferred genome-wide genealogies are accurate enough to faithfully capture the true integer partition distributions. We have attempted to account for the bias introduced by genealogy inference through simulation (see Methods), but we nonetheless expect there to be additional bias in real data that is not captured by our approach. Finally, integer partition distributions from a single population do not contain enough information to infer a fully parameterised demographic model where the *N*_*e*_ of both populations and *m*_*e*_ all vary through time. One potential solution would be to replace an unlabelled lineage in the sample with a labelled lineage that does not receive gene flow, such as a Denisovan (Loya *et al*. 2026; Rogers *et al*. 2026). Increasing the sample size beyond *n* = 5 would be another way to improve estimation. However, rather than developing an estimation method capable of fitting highly complex demographic models, the aim of this work has been to highlight a signal of ancient gene flow that is clear, intuitive and has so far not been used in any population genomic analyses.

## Methods

### Theory

For a panmictic population, the probability of observing integer partition *x* at time *t* given an initial sample of *n* lineages can be calculated as a product of two terms

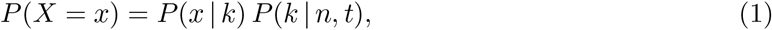

with *k* being the number of lineages in integer partition *x*. The second term is the probability of exactly *k* lineages remaining in the sample at time *t*, which was derived by Tavaré (1984)

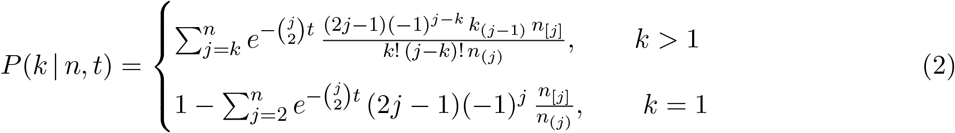

The other term in Equation 1 corresponds to the probability of observing integer partition *x* given that there are *k* lineages. This can be obtained by taking a recursion over the coalescent history from the current *k* up to *n*.

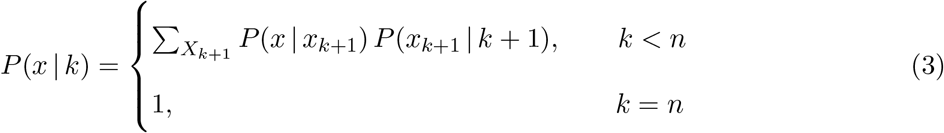

Here, *X*_*k*+1_ is the set of possible integer partitions with *k* + 1 lineages and *P* (*x* | *x*_*k*+1_) is the probability that a coalescence event results in partition *x* given that the previous integer partition was *x*_*k*+1_. The latter can be calculated by enumerating over the possible ways that lineages in *x*_*k*+1_ can coalesce.

Equation 1 can be used to obtain symbolic expressions in terms of *t*. An alternative approach is to define a continuous time Markov Chain where states are integer partitions. As an example, the rate matrix for *n* = 4 under panmixia is shown below with the integer partition states to the right.

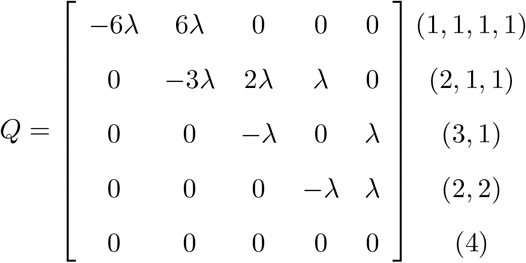

The probability of observing integer partition *x* at time *t* can then be calculated by the matrix exponential *π*_0_ *e*^*Qt*^, with *π*_*o*_ being the initial state vector. This approach can be extended to a structured model by including states where integer partitions are distributed across the two populations. For example, the state (3, 2) expands to [(3, 2), ()], [(3), (2)], [(2), (3)], [(), (3, 2)] in a two population setting. In this model transitions include coalescence at rate *λ* and migration of lineages at rate *m*_*e*_. Details of how the rate matrix is constructed and evaluated can be found in the supplementary python notebook (see Data availability).

### Human genealogies

Ancestral recombination graphs previously estimated using SINGER and POLEGON for the YRI population of the 1000 Genomes Project were downloaded from zenodo (Byrska-Bishop *et al*. 2022; Deng 2025a,b,c). A B-map estimated for the YRI population with predictions at the scale of 1 Mb was also downloaded from zenodo (Buffalo and Kern 2024b,a). Marginal genealogies were sampled from chromosomes every 10 kb using tskit (Kelleher *et al*. 2016; Wong *et al*. 2024) and integer paritions were recorded if the sample position met the following criteria; that positions are at least 5 kb from exonic, telomeric or centromeric regions, that the 1 Mb region encompassing the position has an estimated *B* value between 0.75 and 0.80 and that the nucleotide diversity in the 1 Mb region was well predicted by the *B* value (allowance of *±*10% error). These criteria were designed to minimise the influence of direct and linked selection on downstream analyses while retaining enough loci to accurately estimate integer partitions distributions. For the 10,993 genealogies that passed these criteria, ten subsamples of *n* = 5 from the lineages remaining at 8,000 generations were taken. This subsampling was repeated across ten SINGER MCMC samples to give a total of 100 subsamples per-genealogy. Integer partition distributions between 8,000 and 70,000 generations ago were recorded across 621 discrete time points, with a spacing of 100 generations.

### Model fitting

Models of pulse-admixture and continuous gene flow were fit to empirical integer partitions distributions by maximising the log composite likelihood

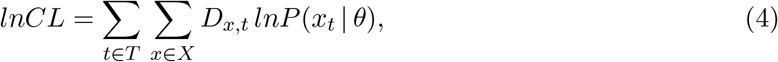

where *T* is the set of discrete time points, *X* is the set of possible integer partitions, *D*_*x,t*_ is the observed frequency of partition *x* and time point *t* and *θ* represents the model parameters. The probability *P* (*x*_*t*_ | *θ*) was calculated by matrix exponentiation across time intervals with piece-wise constant rates. These intervals correspond to ten fixed epochs of equal duration (6,200 generations). Demographic events including pulse admixture, the onset of migration and the merging of populations all introduce additional time points where probability mass is moved between populations or rates of migration change. Maximum composite likelihood estimates for 14 parameters (ten *N*_*e*_*A* values, one *N*_*e*_*B* value, *m*_*e*_ / *α, T*_0_ and *T* 1) were obtained using the Quasi-Newton method in Mathematica (Wolfram-Research 2026). This approach does not accept parameter bounds, but upper bounds of *T*_1_ *≤* 7 *×* 10^4^ and *N*_*e*_*B ≤* 1 *×* 10^5^ were added to the likelihood function to improve convergence and limit parameter space to plausible models.

A 24-epoch panmictic model of population history was estimated from pairwise coalescence times in the human genealogies. These coalescence times were the exact same as those contributing to the integer partition distribution estimates (see above). The coalescent *N*_*e*_ of each epoch was estimated by

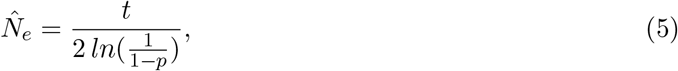

where *t* is the epoch duration and *p* is the proportion of pairs that were uncoalesced at the start of the epoch that coalesce before the epoch end. The epoch duration was defined dynamically so that the number of coalescence events per-epoch were approximately equal.

To account for bias introduced by genealogy inference, we simulated 25 *×* 1 Mb replicates of all inferred population histories using msprime (Baumdicker *et al*. 2021). Rates of recombination and mutation were set to *r* = 1 *×* 10^*−*8^ and *µ* = 1.2 *×* 10^*−*8^, respectively. A multiplicative error profile was calculated for each model by comparing the integer partition distributions from the simulations to those estimated from genealogies after applying SINGER and POLEGON to the simulated VCF file. Specifically, error was defined as *ϵ*_*x,t*_ = *O*_*x,t*_*/E*_*x,t*_, with *O*_*x,t*_ and *E*_*xt*_ being observed and expected integer partitions at time *t*, respectively. We limited error estimates to between 0.1 and 10 and incorporated the error profiles into a second round of model fitting by multiply model predictions by the error profile and normalising so that Σ _*x∈X*_ *P* (*x*_*t*_ | *θ*) = 1.

### Simulations

A panmictic population history with hard selective sweeps was simulated using msprime (Baumdicker *et al*. 2021). The ‘multiple sweeps’ msprime tutorial (https://tskit.dev/msprime/docs/stable/ancestry.html#multiple-sweeps) was adapted so that the waiting times between sweeps were exponentially distributed with expectation 6.4 *×* 10^4^ generations. A total of 25 *×* 1 Mb simulations were performed using a constant recombination rate (*r* = 1 *×* 10^*−*8^) and selection coefficient (*s* = 0.01). A stepping stone model of 30 demes (*N*_*e*_ = 1000, *m*_*e*_ = 1 *×* 10^*−*3^) was simulated in msprime, again with 25 *×* 1 Mb replicates. Sampling locations were randomly chosen and the boundaries parameter was set to True. Integer partitions were recorded from the tree sequences using the same approach as for the human genealogies.

## Data availability

A supplementary python notebook with code for calculating expected integer partition distributions and recording them from tree sequences can be found at the following github repository: https://github.com/A-J-F-Mackintosh/Integer-partition-coalescent. The human integer partitions distributions can be found at the same repository.

## Generative AI use

Anthropic’s Claude (Opus 4.6) was used for assistance writing Mathematica code.

## Acknowledgments

We would like to thank Konrad Lohse, Per Sjödin and José Cerca for helpful feedback on previous versions of this manuscript.

## Supplementary Figures

**Figure S1:**
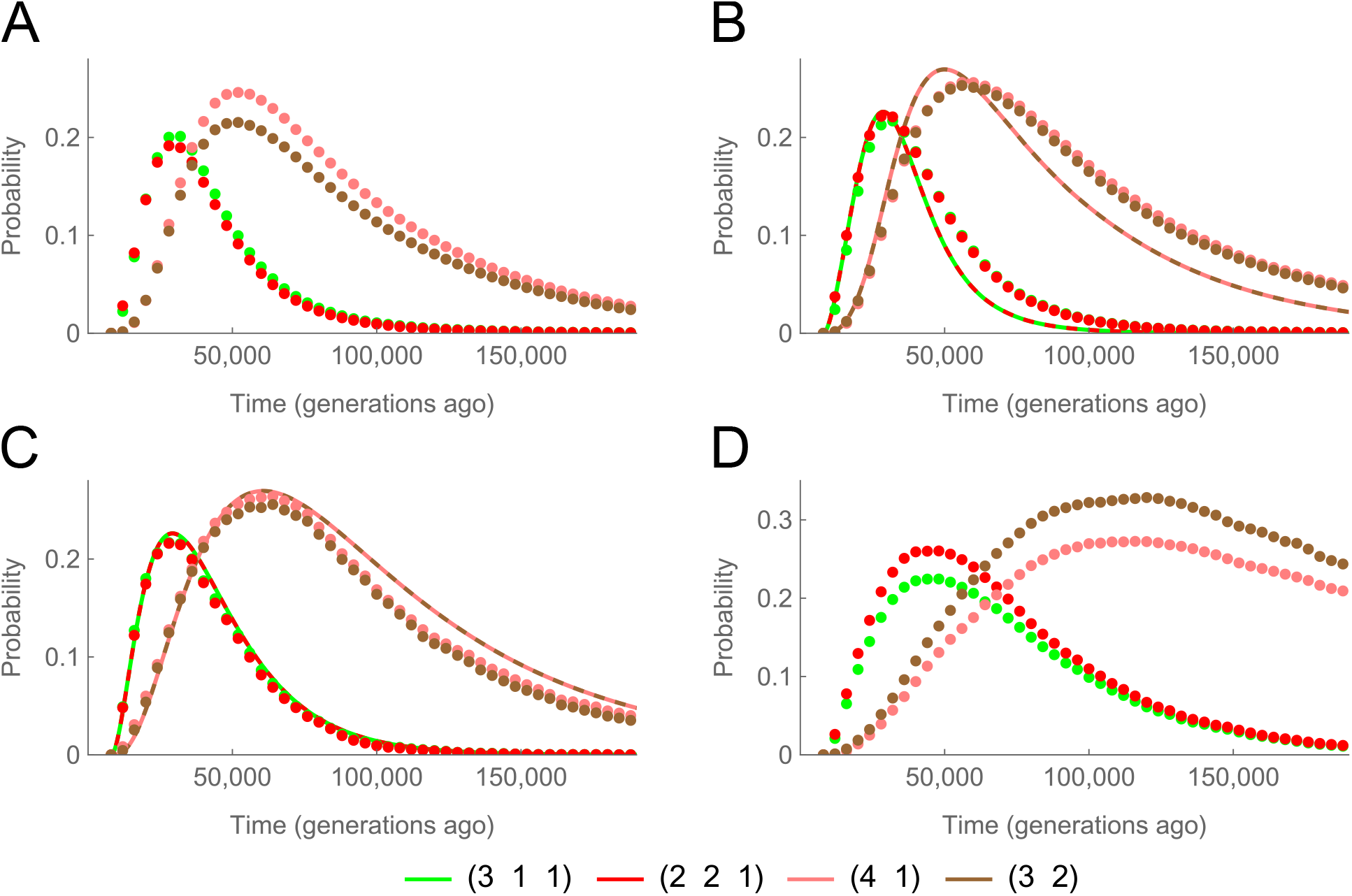
Topologically informative integer partition distributions for data and simulations. **A:** Distributions for human data. **B:** Distributions for the panmictic model estimated from pairwise coalescence times in human data. Lines correspond to the analytic prediction while points are for genealogies sampled by SINGER and POLEGON given a simulated VCF file. **C:** Distributions for a panmictic population with occasional hard selective sweeps. Lines correspond to the analytic prediction without selective sweeps, points correspond to genealogies from simulations that include sweeps. **D:** Distributions for a stepping-stone model of population subdivision estimated by simulation.

